# Individual head models for estimating the TMS-induced electric field in rat brain

**DOI:** 10.1101/2019.12.23.886861

**Authors:** Lari M. Koponen, Matti Stenroos, Jaakko O. Nieminen, Kimmo Jokivarsi, Olli Gröhn, Risto J. Ilmoniemi

## Abstract

In transcranial magnetic stimulation (TMS), the initial cortical activation due to stimulation is determined by the state of the brain and the magnitude, waveform, and direction of the induced electric field (E-field) in the cortex. The E-field distribution depends on the conductivity geometry of the head. The effects of deviations from a spherically symmetric conductivity profile have been studied in detail in humans. In small mammals, such as rats, these effects are more pronounced due to their smaller and less spherical heads. In this study, we describe a simple method for building individual realistically shaped head models for rats from high-resolution X-ray tomography images. We computed the TMS-induced E-field with the boundary element method and assessed the effect of head-model simplifications on the estimated E-field. The deviations from spherical symmetry have large, non-trivial effects on the E-field distribution: in some cases, even the direction of the E-field in the cortex cannot be reliably predicted by the coil orientation unless these deviations are properly considered.

## Introduction

In transcranial magnetic stimulation (TMS), a pulsed magnetic field induces an electric field (E-field) in the brain, causing action potentials in the targeted cortical region. In a typical TMS pulse, the magnetic field is increased from 0 to 1–2 tesla in less than 100 μs, which induces an E-field of the order of 100 V/m. TMS has established clinical applications in, e.g., presurgical mapping of brain functions^1–4^ and treatment of drug-resistant major depression^5–7^. Promising clinical applications are emerging, such as stroke rehabilitation^8–10^ and treatment of chronic pain^11,12^.

The TMS-induced E-field accumulates charge at cell membranes, causing their local de- or hyperpolarisation^13^. This can happen in several ways: The E-field gradient polarises long straight sections of an axon, the E-field polarises terminations and sharp turns of an axon, or a discontinuity in the E-field due to different tissue properties polarises an axon passing through the tissue interface. The last two are likely the strongest mechanisms^14^. In all three cases, the polarisation depends on the direction of the induced E-field and is proportional to its magnitude^14^.

Optimal interpretation of data obtained by combining TMS with functional brain imaging requires knowledge of the induced E-field distribution. For example, different E-field orientations even in the same target location recruit different neuronal populations^15^. Computing the E-field requires a model of the electrical conductivity distribution of the head, often in the form of a volume conductor model (VCM). Sometimes a simple VCM is entirely adequate: For example, for human TMS with the coil near the vertex, the location of the resulting E-field maximum can be computed quite accurately with a spherically symmetric VCM fitted to the local radius of curvature^16^. However, even for human TMS, such a spherical model can be inadequate when stimulating the frontal or occipital lobes: The E-field-magnitude prediction can be off by tens of percent—even when only considering the error relative to the E-field-magnitude prediction in the motor cortex, i.e., the prediction error relevant to a typical experiment where the desired stimulation intensity is proportional to individual motor threshold^16^. For rodent TMS, the required level of detail in the VCM has not been studied.

There is increasing interest to use TMS in rodents due to a myriad of genetically modified animals and disease models available. As more invasive recording techniques can be used in rodents than in humans, and as brain tissue is available for histological and molecular analysis, rodent TMS studies are valuable, e.g., in studying brain plasticity caused by repetitive TMS^17^ and TMS-induced disease modification^18^. Rodent studies allow also assessing, e.g., gene expression following repetitive TMS^19,20^. In previous experimental TMS studies involving rats, the head has been approximated with a body-shaped volume of uniform conductivity^21^, with concentric ellipsoids^22^, or with a spherical model^23^. In addition, the infinite homogeneous conductor model has been used^24^; for a rat, the latter model overestimates the induced E-field by a factor between five and eight^25^ as it fails to capture the reduction in stimulation efficiency for coils much larger than the head^26^. In such cases of not having a finite VCM and overall in cases, where a VCM is not used, the TMS experimenters often target the stimulation using a line-of-sight model (this is called line navigation), in which the E-field maximum is assumed to be directly below the coil centre and E-field orientation is assumed to correspond to coil orientation^24^. For humans, subject-specific head models are derived from magnetic resonance images using pipelines that combine various pieces of software, such as FreeSurfer^27^, FSL^28^, and SPM^29^. For the rat head, we are not aware of any proposed recipe for combining software algorithms to create subject-specific realistic head models. Rather, modelling studies often use post-mortem anatomical atlas data for their animal models^26,30^.

Here, we describe a simple processing pipeline for generating individual head models from micro-scale computed tomography (μCT) data and computing the E-field using reciprocity and the boundary element method (BEM). With the resulting efficient procedure, we computed TMS-induced E-fields in a rat head using VCMs with varying level of detail. We compare four levels of detail: line navigation, a spherically symmetric head model fitted to the inner skull surface, a realistically shaped single-compartment (1C) model that follows the body outline of the rat similarly to the model used by Salvador and Miranda^21^, and two-compartment (2C) models that contain the body outline and the skull. As a reference model, we use a high-density model which further includes the spine and eyes.

## Results

We built five different head models for one adult male Wistar rat—a spherical model, a 1C model, two 2C models (with and without spine, respectively), and a high-resolution reference 3C model (with the spine and eyes)—and computed the TMS-induced E-field for 35 different coil positions shown in Fig. 1. These positions span a 10 mm × 15 mm region on the scalp, covering the superior parts of the cortex. For each position, we studied all tangential coil orientations with a 10° step size, resulting in 630 unique coil placements. Compared to the reference model, the cortical E-field in the 2C model with spine had a relative error (RE) of 7.6±1.6% over all coil positions and orientations (mean ± standard deviation) and 4.7±2.4% in the region of the strongest fields (i.e., where the E-field energy density exceeded 50% of its maximum, see Methods); omitting the spine, the error increased to 9.6±2.4% (5.1±2.2% in the region of the strongest fields). Both of these errors were, however, far smaller than the errors arising from omitting the skull: the 1C model had an RE of 64±10% (41.1±4.6% in the region of the strongest fields), making it worse than the spherical model (RE 45.0±2.5%, and 35.2±7.0% in the region of the strongest fields). The RE is sensitive to all types of errors in the E-field prediction, including its overall magnitude. Errors in the overall magnitude, however, have a limited effect on typical TMS experiments, where the stimulation intensity is normalised to an experimentally determined motor threshold. Omitting the general magnitude, the correlation errors (CCE) of the E-field patterns were 0.53±0.23%, 0.73±0.33%, 16.2±2.9%, and 17.5±2.8%, respectively. For the region of the strongest fields, the correlation errors increased to 0.9±1.0%, 1.1±1.3%, 54±25%, and 35±25%, respectively. Further omitting even local magnitudes, the mean angular errors in the direction of the cortical E-field were 3.5±0.6°, 4.6±0.9°, 26.8±4.6°, and 25.1±1.8°, respectively. For the region of the strongest fields, the mean angular errors were reduced to 1.6±1.2°, 1.5±1.0°, 10.8±4.4°, and 13.7±3.6°, respectively. As the difference between the two 2C models in the region of strongest fields was relatively small with all these measures, we will for clarity omit the 2C model with spine from Fig. 2.

**Figure 1:**
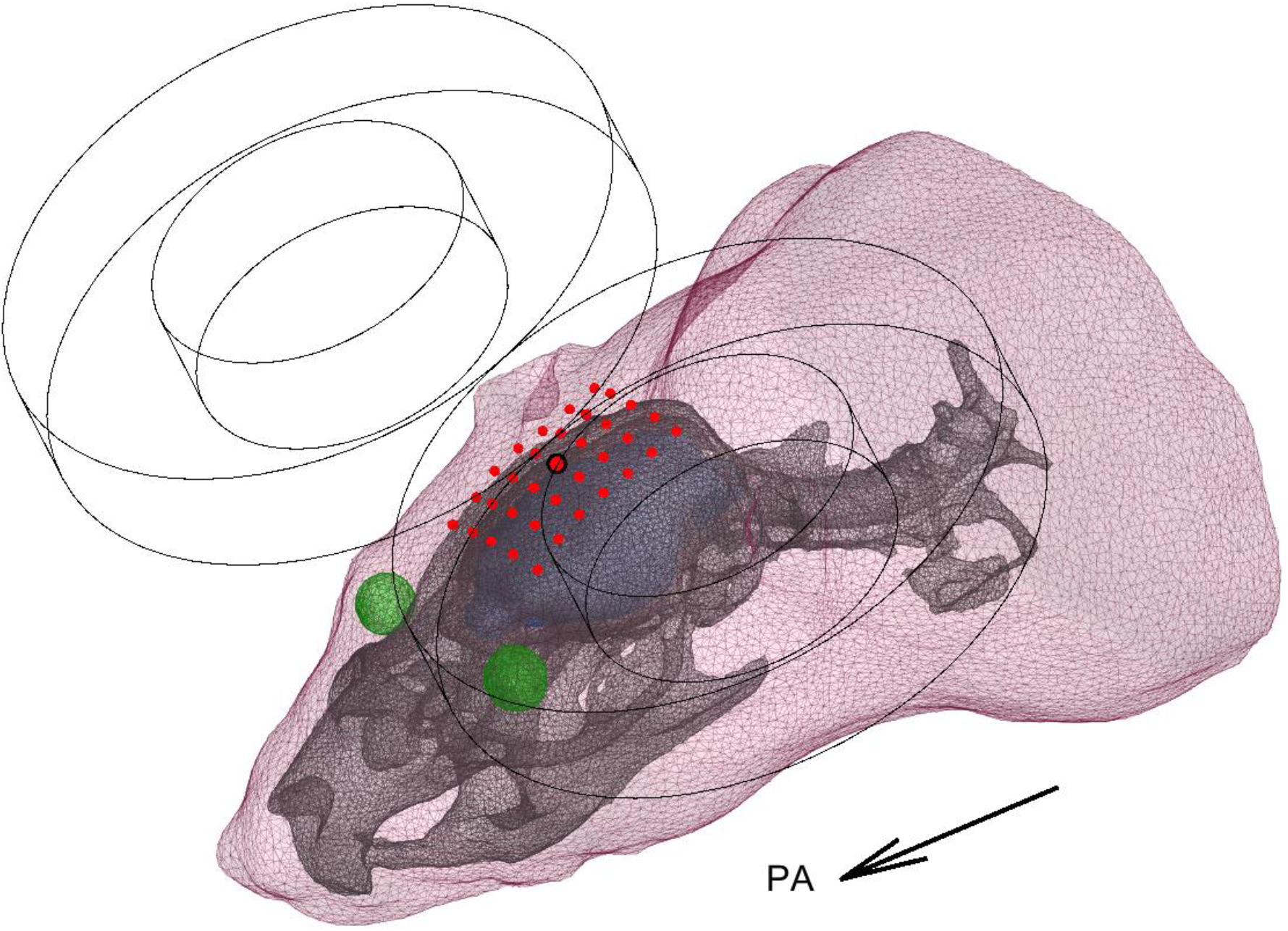
A three-compartment BEM model of the rat head. The compartments are defined by the body outline of the front half of the rat, the skull, and the eyes. The induced E-field was computed on the blue surface, which corresponds to the estimated surface of the brain and the brainstem. The red points indicate the tested locations of the coil centre, with the central point highlighted. Here, an example coil is shown in the posterior–anterior (PA) orientation at the central location.

**Figure 2:**
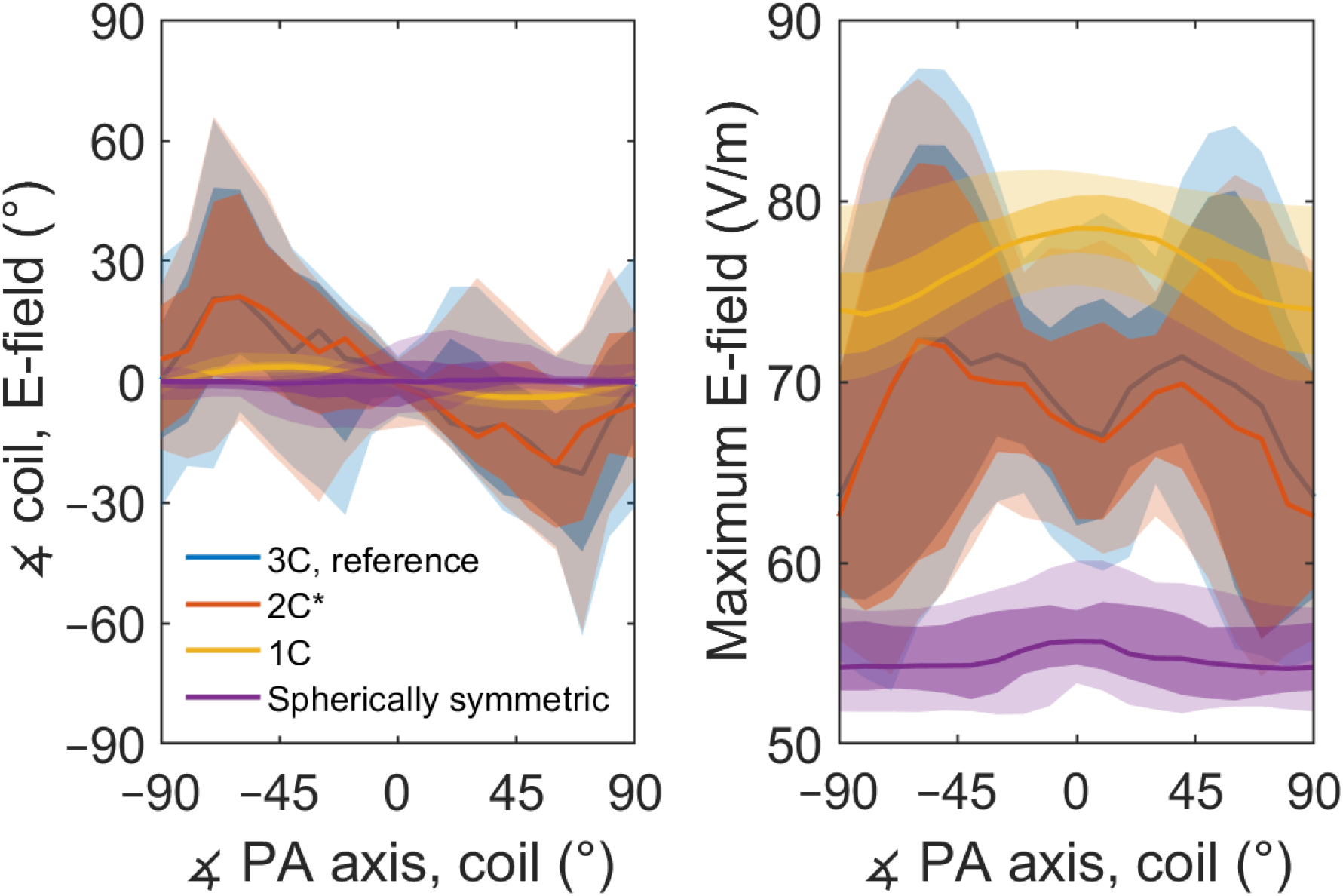
Model differences depend on the coil orientation. *Left:* The difference between the coil orientation and the direction of the E-field at the E-field maximum for all coil locations, at different angles with respect to the posterior–anterior (PA) orientation. To get a directed 1-dimensional angle, the E-field was projected to the plane defined by the bottom of the coil. *Right:* The distribution of the peak E-field over all tested coil locations at different coil orientations. The lighter colours cover percentiles from 2.3% to 97.7%, the darker colours percentiles from 16% to 84%, corresponding to ±2 and ±1 standard deviations for normally distributed data, respectively. The solid lines represent the median values. 2C* refers to the spineless 2C model, for model geometries, see Fig. 4.

In realistic head geometry, the E-field magnitude depended on the coil orientation. Starting from the posterior–anterior (PA) orientation shown in Fig. 1, the peak E-field magnitude first increased when the coil was rotated to either direction, but then dropped sharply when the coil was close to ±90° from the PA direction (Fig. 2). Because of this behaviour, the E-field orientation near the E-field maximum was closer to the PA direction than could be expected from the coil orientation. That is, in the left panel of Fig. 2, the direction of the error has opposite sign to the coil orientation. With both the reference model and the 2C models, the error increased until an angle of about ±70°, at which point the discrepancy between the coil and the E-field direction varied between –10 and 70° depending on the coil location. The 1C model captured only a fraction of this effect; in the 1C model the stimulation direction was on average closer to the PA direction than could be expected from the coil orientation, but the difference was only a few degrees instead of up to several tens of degrees. In the spherically symmetric model, the median orientation error over all locations was close to zero. There was, however, a coil-location-dependent orientation error, which resulted from the coil locations following the surface of the scalp and not that of the spherical geometry. The orientation-dependence of the field magnitude was likely due to the oblong shape of the rat head and the cranial compartment, as almost no such drop occurred in the spherical model.

Defining the true stimulation target as the location of the maximum of the induced E-field in the cortex in the realistic head geometry, the true target was systematically more posterior and medial than expected from line navigation, the spherical head model, or the 1C head model. The error depended on the coil orientation and location. The median error for line navigation was 7.3 mm (for 5% of the coil placements the error was greater than 12.7 mm); for the spherical head model and the 1C model, the errors were 6.3 mm (95%: 12.2 mm) and 7.2 mm (95%: 12.0 mm), respectively; whereas the 2C models with and without spine had median errors of just 0.3 mm (95%: 6.0 mm) and 0.4 mm (95%: 7.2 mm), respectively. A cross comparison of the prediction differences between the models is shown in Fig. 3. Line navigation, the spherical head model, and the 1C head model produced similar, similarly incorrect, predictions. The large systematic errors in the predicted stimulation location highlight the importance of an adequate model. In addition to the systematic errors, the data underlying Figs. 2 and 3 contain further several coil-placement-dependent outliers that could not be predicted without modelling the skull. These largest prediction errors occurred when either the coil orientation was close to the PA orientation and the true stimulation maximum was in the brainstem, as seen in, e.g., Fig. 4, or the coil orientation was such that a large part of the windings passed directly above either eye of the rat and the true stimulation maximum was close to the hole in the skull behind that eye irrespective of the actual coil position. In the first case, the E-field maximum corresponded to the location in which the brainstem passes through the back of the skull. That is, a larger proportion of the induced currents flowed through the path of least resistance, and consequently an E-field maximum arose in the proximity of the hole in the skull. In the second case, a similar phenomenon occurred with the hole behind the eye.

**Figure 3:**
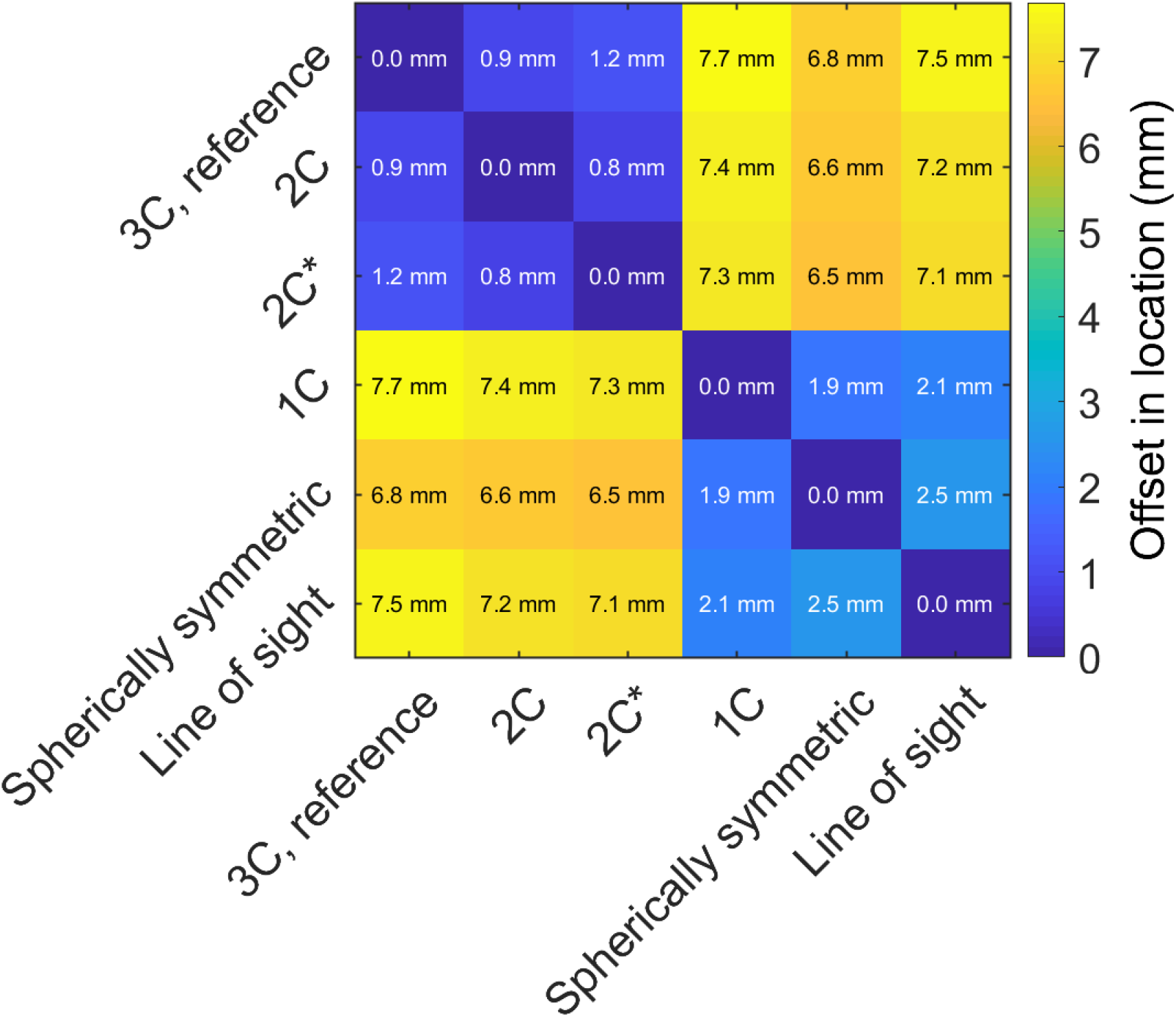
The mean distance between the predicted locations of the E-field maximum between different methods. Varying the level of detail in the skull model had little effect, but omitting the skull geometry resulted in a large, systematic error in the location of the E-field maximum. The true stimulation target was typically more posterior and medial than that predicted by the 1C, spherically symmetric, or line-of-sight models. 2C* refers to the spineless 2C model, for model geometries, see Fig. 4.

**Figure 4:**
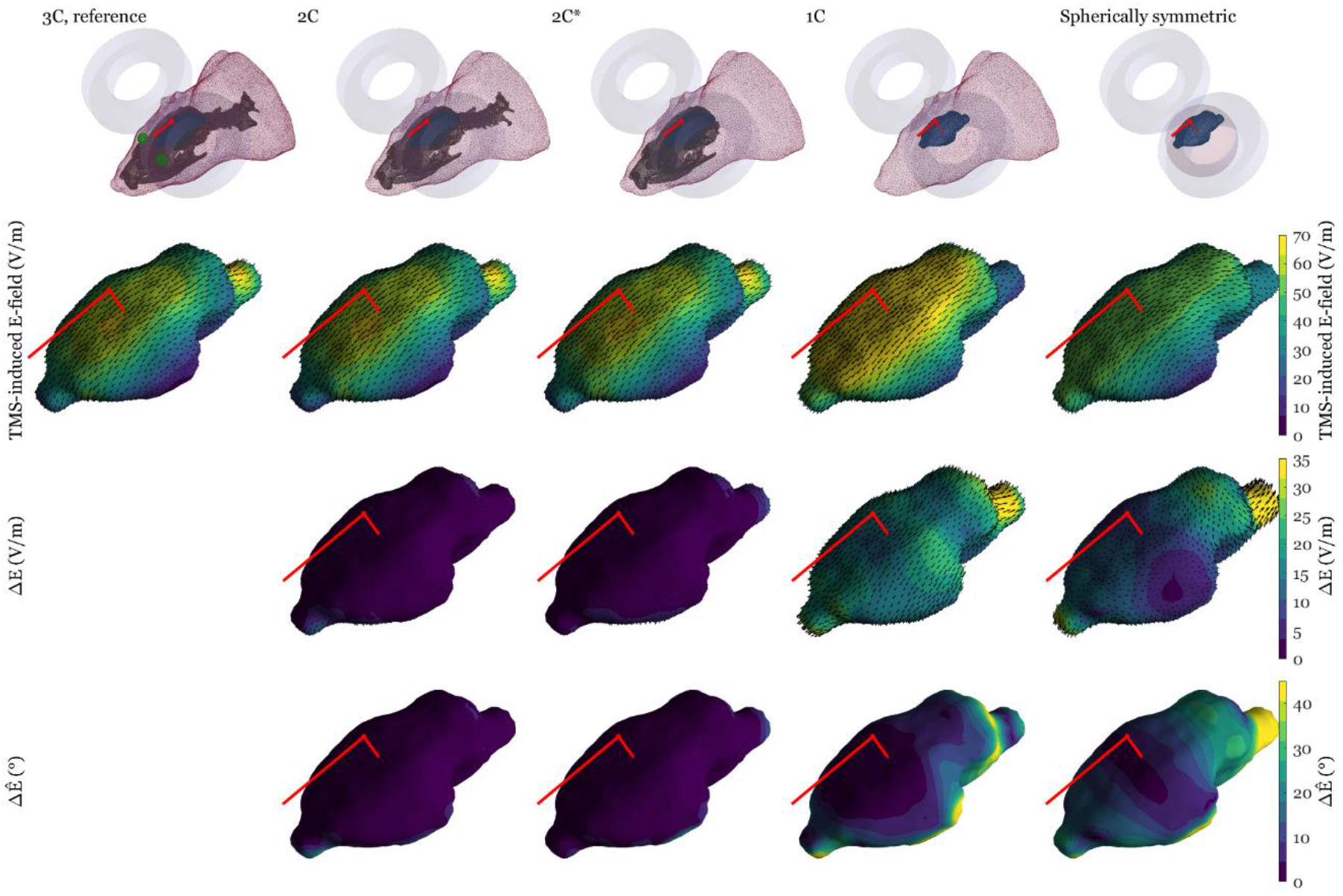
The TMS-induced E-field in the cortex in an example case where the models that omit the skull geometry fail to predict the true stimulus location. From left to right: The high-resolution reference model (1^st^ column), the 2C model with spine (2^nd^ column), the 2C model omitting spine (3^rd^ column), the 1C model (4^th^ column), and the spherical model (5^th^ column). The first row depicts the model geometries; the second row shows the norm of the induced E-field and its direction (dark red arrows); the third row displays the norm and direction of the difference in the E-field compared to the reference model (ΔE); the fourth row shows the difference in direction compared to the reference model (ΔÊ). The bright red lines indicate the coil orientation and the line of sight from the coil centre.

## Discussion

We described a simple method for producing subject-specific realistic-head-geometry BEM models for computing the TMS-induced E-field in small rodents. Compared to line navigation, a subject-specific realistic model allows better estimation of the location and extent of the stimulation. Compared to even a subject-specific spherical model, the realistic model further allows better estimation of the stimulation dose. In addition, we demonstrated that the features of the skull geometry—especially the holes in the skull—have a large effect on the E-field distribution. As the described method can model these holes in the skull, it can accurately model two most influential parts of the head geometry, the head shape and the skull surface in small rodents and other animals with relatively large holes in their skulls. Due to the difficult extraction of soft-tissue boundaries from X-ray tomography images, the differences between different soft tissues were not modelled.

We compared our 2C model with previously used simpler ones and demonstrated that both the 1C model of the rat and the spherical approximation of its skull failed to capture the characteristic behaviour of the TMS-induced E-field in the rat brain. Further, the spherical approximation caused a much larger error in the E-field calculation in the rat than what had previously been observed in humans^16^. This is most likely due to a combination of two things: a less spherical head shape and, relative to the head, a much larger coil in rat TMS. In realistic head geometry, the E-field maximum is generally not directly below the coil and the peak amplitude of the E-field depends strongly on the coil orientation. The 1C model manages to capture some of these effects, qualitatively, but underestimates their strength and is unable to see the E-field focusing caused by holes in the skull. As this focusing effect seems to cause the largest differences between the models, the next logical improvement to our 2C head model would be adding more detail to these regions (such as the eyes in the reference model). Another possibility is to add further tissue compartments spanning through the holes in the skull (which would be the case with cerebrospinal fluid), with, for example, a BEM solver that supports the use of junctioned geometry^31^.

The main source for numerical uncertainty in surface-based models arises from the discretisation of the modelled surfaces. We observed only a small difference (correlation error about 1%) due to halving the mesh density of the head model. In addition, the coil model resolution had only a small effect: To test that the coil model was sufficiently accurate, we substituted our high-resolution coil model with a simplified low-resolution model, which also assumed the 9-mm-tall windings thin. This underestimated the E-field by 3% in all volume conductor models, but produced otherwise essentially indistinguishable distributions with, e.g., less than 0.5° angular error in each model.

## Conclusion

The small and far-from-spherical head of the rat emphasises the effects of conductivity boundaries on TMS-induced E-field distributions compared to those seen in the human head. Correct determination of the stimulus location and orientation requires considering these effects, when TMS experiments with small animals are designed.

## Acknowledgements

This research was supported by the Finnish Cultural Foundation, the Jane and Aatos Erkko Foundation, and the Academy of Finland (Decisions No. 255347, 265680, 283105, and 294625).

## Methods

### Head model generation

In a realistically shaped magnetoencephalography (MEG) or TMS forward model, the inner-skull surface has the largest impact on the volume current flow. To accurately represent this surface, we built a model based on a high-resolution X-ray tomography data (Flex CT, GMI, Northridge, CA, USA) of the head of a male Wistar rat (weight 617 g). The animal imaging procedures were approved by the University of Eastern Finland animal care committee and performed in accordance with their regulations and with the guidelines of the European Community Council Directives 2010/63/EU. The imaging was done using isotropic 0.17-mm voxels (cone beam acquisition, 256 projections, 2-by-2 binning, 512 × 512 × 512 image matrix, 60 kVp tube voltage, 450 μA tube current). The surface meshes that represent conductivity boundaries were extracted by segmenting the image into air, body, and bone. The imaging modality provided good contrast between these three, allowing simple but robust segmentation: We set thresholds halfway between the two main peaks in the intensity histogram (air and sum of all soft tissues, respectively; with our threshold at approximately –500 Hounsfield units (HU)) and after the end of the soft-tissue peak at approximately 500 HU. The segmented body was smoothed by computing its morphological closure with a 2-mm-radius spherical kernel (in addition to smoothing the surface, this operation filled small channels in the nose and ears, considerably reducing the number of vertices needed to model the body outline), and the skull was smoothed similarly with a 1-mm-radius spherical kernel. Finally, two boundary surfaces were extracted from volumetric tetrahedral meshes generated with “vol2mesh” function (iso2mesh, iso2mesh.sourceforge.net)^32^. The resulting high-resolution meshes for the reference model are shown in Fig. 1. The body mesh consists of 13037 vertices and 26070 faces (average edge length 0.95 mm, with a standard deviation of 0.12 mm), and the skull–spine mesh consists of 24318 vertices and 48804 faces, with an average edge length of 0.54 mm (standard deviation 0.12 mm). As the rat skull contains several large holes anterior, posterior, and inferior to the brain volume, one cannot model it using closed inner- and outer-skull surfaces as is conveniently done with human skulls. This poses a problem, as common meshing tools and BEM solvers assume that each boundary surface is closed and that the conductivity contrast across each boundary is constant. We overcame this problem by requiring that the intracranial and body conductivities are equal—a common assumption in conductivity parameters also in human studies. Then, we connected the inner- and outermost volumes which allowed interpreting the three-compartment model as a two-compartment (2C) model where we have one closed skull surface (with at least one hole) floating inside the body surface. With this approach, we lose the ability to simply have different conductivity values for the intracranial volume and the rest of the body but gain the ability to easily model any number of holes in the skull. From computational point of view, further conductivity details can be added by adding further geometric entities inside the body surface: For the reference model, we added the eyes, which could also be extracted from the μCT data. Unlike with the obvious high-contrast interfaces from air to body and from body to skull, the process for extracting the eyes could not be automated, as the signal-to-noise ratio between the eyes and their surroundings was far below 1. We localized the eyes from a 3-dimensional visualization of the body surface, surrounded them with 8.33-mm cubic bounding boxes, and extracted them from their respective bounding boxes by applying median filtering (7 × 7 × 7 kernel, effective resolution 1.2 mm), and compensated for the slight field inhomogeneity in the μCT data by manually selecting separate isolevels to extract either eye. The mesh for eyes has 479 vertices and 950 elements, which brought the total vertex count for the reference model to 37834 and the total element count to 75824. For the 2C model, the skull–spine was independently meshed with a mean edge length of 0.67 mm, resulting in 15238 vertices and 30636 elements. The spineless skull was meshed with the same resolution, which resulted in 12426 vertices and 24964 elements. Finally, the two 2C models and the 1C model used a body mesh with a mean edge length of 1.4 mm (5924 vertices and 11844 elements).

The extent of the brain inside the intracranial cavity was estimated visually, and the corresponding volume was filled with manual voxel painting with GNU Image Manipulation Program (www.gimp.org). The E-field was computed on a surface spanned in the outermost part of this brain volume, at the depth of at least 1 mm from the skull. This distance was chosen to ensure the numerical stability of the solver. The surface estimation was done using a volumetric mask (i.e., subtracting a 1-mm dilated skull mask from the brain mask) and then triangulating the boundary of this modified brain volume.

### Spherically symmetric head model

There is a relatively large region of hole-free skull above the centre of the rat head, where we fitted a sphere based on the local inner-skull surface; the centre of the fitted sphere (origin) was 14.4 mm from the local inner-skull surface (Fig. 4), the maximum distance from the cortex to the origin being 15.5 mm. The shortest distance between the scalp at the coil locations and the origin was 15.4 mm. We could thus fit a sphere that was fully between the brain volume and the nearest part of coil windings, meaning that the E-field in the whole brain could be computed using this spherical model (although the spherical head model does not depend on the radial conductivity profile and thus on the head size, the model is only valid for sources at radii smaller than the smallest distance from the origin to the coil^33^).

We needed to calculate the induced E-field everywhere in the cortex for different coil positions and, ideally, would have liked to fit local spheres for all those positions. The shape of the rat head made this re-fitting problematic, as, for most non-central positions, the “other end” of the brain would no longer have been inside such a sphere, and the model would have been both physically invalid and mathematically wrong for those remote regions. Here we overcame this problem by interpreting the centrally fitted sphere as the global spherical approximation of the head, as it nicely covered the whole brain, and was in size similar to, e.g., the spherical model used in^23^. We computed the E-field in the spherical model with the analytical closed-form solution^33^.

### Boundary element method

The TMS forward problem being reciprocal to the MEG forward problem^34^, the TMS-induced E-field can be obtained by computing the MEG forward problem with the TMS-coil as the pick-up coil^35^. The boundary element method is increasingly popular for solving this MEG forward problem. BEM is tailored for problems where the geometry can be described by a few piecewise homogeneous isotropic regions, such as the brain, cerebrospinal fluid, skull, and scalp (or in our case, mean body tissue). Key benefits of using a BEM instead of a finite element model or some other volume-based model are its lower memory and computation-time requirement and the ease of segmentation, meshing and visualisation, as only the extracted conductivity boundaries need to be meshed and discretisation is performed only on these boundary surfaces. Consequently, a typical high-resolution 3S model consists of less than 10000 vertices and can be solved for a large number of source and field parameters with a regular workstation computer in minutes^36^. With further optimizations, the E-field computation for each coil position for models can be computed in real time, in just a few tens of milliseconds^37^.

In this study, the TMS-induced E-field in the 1C and the two 2C models were computed using a linear-collocation (LC) BEM implementation of the Helsinki BEM framework^36^, further developed from the Helsinki BEM library^39^. The conductivity of the combined intracranial–body compartment was set to 0.33 S/m and the skull conductivity to 6.6 mS/m (the ratio between the two conductivities being 1/50). To confirm that a simple and fast LC solver was valid with the complicated skull mesh that contains thin structures, the high-resolution 3C reference model was built using a much slower, but generally more accurate and robust, linear Galerkin (LG) solver^40,41^. In the reference model, the conductivity value for the eyes was set to 1.5 S/m.

### Coil model

We modelled the MagVenture MC-B35 Butterfly Coil (MagVenture A/S, www.magventure.com), which is a small figure-of-eight coil (for dimensions, see Table 1), with a set of magnetic dipoles whose locations and weights are those of the integration quadrature points for computing the magnetic flux integral for MEG^42^. We built two coil models for the 9-mm-thick coil windings, one with a total of 2568 dipoles in three identical layers (at 1.5, 4.5, and 7.5 mm from the lowermost wire surface) and one with lower resolution, with just one layer (at 4.5 mm) with 136 dipoles.

**Table 1:** MagVenture MC-B35 Butterfly Coil dimensions according to manufacturer’s website (February 24, 2016) and private conversation with the manufacturer’s representative.

|  |  |
| --- | --- |
| Inner winding diameter | 24 mm |
| Outer winding diameter | 47 mm |
| Wire thickness | 9 mm |
| Insulation thickness between wires and the coil<br>bottom | 2 mm |
| Number of turns | 2 wings, with 3 layers of 4 turns each |

### Error metrics

We used three distinct metrics for evaluating the differences between the models similar to ^37^. First, we computed the relative error

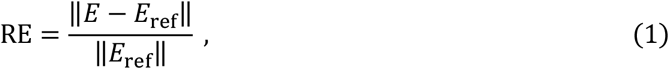

where *E* and *E*_ref_ have been pooled into 3*N* × 1 vectors, where *N* is the number of evaluation points in the cortex (*N* = 1833), describing the three Cartesian components of the induced E-field distribution of the tested model and the reference model (3C), respectively. This measure is sensitive to errors in both magnitude and direction of the induced E-field. Second, we computed the correlation error (CCE)

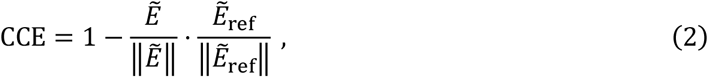

where *Ẽ* and *Ẽ*_ref_ have first been de-meaned component-wise across field points and then pooled into 3*N* × 1 vectors. This measure is primarily sensitive to overall topographical differences of the E-field; it, however, omits the possible constant-factor difference in the E-field magnitude between different models. Third, we computed the mean of the angular errors between the E-fields at respective locations. This measure is sensitive only to errors in the field direction. These three error metrics were computed both for all points and only for those points where the E-field energy density was over 50% of its maximum (i.e., the E-field magnitude was more than √0.5 times the peak magnitude for that coil location).

## Data availability

Data available on request from the authors.

## Author contributions

L.M.K., J.O.N., O.G., and R.J.I. conceived and designed the research. K.J. acquired the X-ray tomography data, and L.M.K. segmented the body, skull and eyes from the data. L.M.K. and M.S. designed the simulations. M.S. designed and implemented the forward field solver and L.M.K. ran the simulations. L.M.K., M.S., and J.O.N. wrote the initial version of the manuscript. All authors participated in finalising the manuscript.

## Competing interests

The authors declare no competing interests.

